# Relationship between reactive astrocytes, by [^18^F]SMBT-1 imaging, with amyloid-beta, tau, glucose metabolism, and microgliosis in mouse models of Alzheimer’s disease

**DOI:** 10.1101/2023.08.21.554163

**Authors:** Yanyan Kong, Cinzia A. Maschio, Xuefeng Shi, Bolin Yao, Fang Xie, Chuantao Zuo, Uwe Konietzko, Kuangyu Shi, Axel Rominger, Jianfei Xiao, Qi Huang, Roger M. Nitsch, Yihui Guan, Ruiqing Ni

**Author notes:** Corresponding author: Yihui Guan, Ruiqing Ni.

## Abstract

**Purpose:** Reactive astrocytes play an important role in the development of Alzheimer’s disease (AD). Here, we aim to investigate the temporospatial relationship between reactive astrocytes, tau and amyloid-β, glucose metabolism, and microgliosis by using multitracer imaging in AD transgenic mouse models.

**Methods:** Positron emission tomography (PET) imaging with [^18^F]SMBT-1 (monoamine oxidase-B), [^18^F]florbetapir (Aβ), [^18^F]PM-PBB3 (tau), [^18^F]FDG, and [^18^F]DPA-714 (translocator protein) was carried out in 5- and 10-month-old APP/PS1, 11-month-old 3×Tg mice, and aged-matched wild-type mice. The brain regional referenced standard uptake value (SUVR) was computed with the cerebellum as the reference region. Immunofluorescence staining was performed in mouse brain tissue slices.

**Results:** [^18^F]SMBT-1 and [^18^F]florbetapir SUVRs were higher in the cortex and hippocampus of 10-month-old APP/PS1 mice than in 5-month-old APP/PS1 mice and wild-type mice. Reduced [^18^F]FDG SUVR was observed in the thalamus and midbrain of 5-month-old APP/PS1 mice compared to wild-type mice. No significant difference in brain regional [^18^F]DPA-714 SUVR was observed in 5- and 10-month-old APP/PS1 mice compared to wild-type mice. No significant difference in the SUVRs of any tracers was observed in 11-month-old 3×Tg mice compared to age-matched wild-type mice. A positive correlation between the SUVRs of [^18^F]SMBT-1 and [^18^F]DPA-714 in the cortex was observed. Immunostaining validated the distribution of MAO-B and TSPO, amyloid and tau inclusions in brain tissue from 10-month-old APP/PS1 mice and limited changes in 11-month-old 3×Tg mice.

**Conclusion:** The findings provide in vivo evidence for reactive astrocytes along with amyloid plaque and tau deposition preceding microgliosis in animal models of AD pathologies.

## 1 Introduction

Alzheimer’s disease (AD) is pathologically characterized by abnormal accumulation of amyloid-beta (Aβ), tau tangles, reactive astrocytes, microgliosis and neuronal loss. Astrocytes are the most abundant glial cell population in the brain and play an important role in maintaining synaptic homeostasis by regulating synapse function, calcium signalling, and brain metabolism [1]. Reactive astrocytes are involved early in the pathophysiology of AD [2–4]. Reactive astrocytes measured by cerebrospinal fluid levels of glial fibrillary acidic protein (GFAP) have been shown to mediate the effect of Aβ on tau and drive downstream neurodegeneration and cognitive impairment in patients with AD [5] and preclinical AD. Postmortem studies of AD brains have demonstrated abundant reactive astrocytes and microglia around Aβ plaques and tangles [6–8]. Monoamine oxidase B (MAO-B) levels are increased not only in astrocytes but also in pyramidal neurons in the AD brain. Reactive astrocytes and MAO-B have emerged as treatment targets for neurodegenerative diseases [9].

Several positron emission tomography (PET) tracers for reactive astrocytes have been developed, including the MAO-B tracers [^11^C]deuterium-L-deprenyl (DED) and [^18^F]F-DED [10–12], the novel tracer [^18^F]SMBT-1, the mitochondrial imidazoline2 binding site (I_2_BS) tracer [^11^C]BU99008 [5], [^11^C]acetate [13] and the thyroid hormone transporter OATP1C1 [^18^F]sulforhodamine-101 [14]. In vivo [^11^C]DED has demonstrated divergent longitudinal changes in reactive astrocytes and amyloid in patients with autosomal-dominant [15] and prodromal AD [16, 17]. Higher brain [^18^F]SMBT-1 binding in Aβ+ than in Aβ-nondemented controls has been observed and is associated with Aβ accumulation at the preclinical stage of AD [18–22]. In animal models (APPswe, PS2APP, APPArcSwe), reactive astrocytes measured by using [^11^C]DED and [^18^F]F-DED precede the increase in the amyloid-PET signal [10–12].

The aim of the current study was to evaluate the distribution of the novel tracer [^18^F]SMBT-1 in two mouse models of AD (APP/PS1, 3×Tg). Using a multitracer approach with [^18^F]SMBT-1, [^18^F]florbetapir, [^18^F]PM- PBB3 (florzolotau, APN-1607), [^18^F]fluorodeoxyglucose (FDG), and [^18^F]DPA-714 (translocator protein (TSPO)), we assessed the temporospatial relationship of reactive astrocytes with Aβ, tau, glucose metabolism, and microgliosis. We hypothesized that reactive astrocytes are an early event associated with tau and Aβ accumulation. To our knowledge, this is the first in vivo imaging study using [^18^F]SMBT-1 in animal models of AD, along with tracers for the aforementioned other pathophysiologies.

## 2 Methods

### 2.1 Animal models

The animal models used in the study are summarized in **Table 1**. 3×Tg mice [B6;129- Psen1tm1MpmTg(APPSwe, tauP301L)1Lfa/Mmjax] aged 11 months [23], and APP/PS1 mice [B6. Cg- Tg(APPswe,PSEN1dE9)85Dbo/Mmjax] mice overexpressing the human APP695 transgene (Swedish (K670N/M671L)) and PSEN1 mutations [24] aged 5 and 10 months were used (Jax Laboratory, USA). Wild-type C57BL6 mice were obtained from Charles River Germany and Cavins Laboratory Animal Co. Ltd. of Changzhou. Mice were housed in ventilated cages inside a temperature-controlled room under a 12-h dark/light cycle. Pelleted food (3437PXL15, CARGILL) and water were provided ad libitum. Paper tissue and red Tecniplast mouse house® (Tecniplast, Italy) shelters were placed in cages for environmental enrichment.

**Table 1.** Information on the animal models used in the study.

| **Animal model** | **Age (month)** | **[<sup>18</sup>F]SMBT-1** | **[<sup>18</sup>F]DPA-714** | **[<sup>18</sup>F]PM-PBB3** | **[<sup>18</sup>F]AV45** | **[<sup>18</sup>F]FDG** | **[<sup>11</sup>C]PIB** |
| --- | --- | --- | --- | --- | --- | --- | --- |
| APP/PS1 | 5 | 3 M | 3 M | 3 M | 7 M | 3 M |  |
|  | 10 | 3 M | 3 M |  | 6 M | 3 M |  |
| 3×Tg | 11 | 8 M | 3 M | 5 M | 3 M | 3 M | 2F/2 M |
| Wildtype | 5 | 6 M | 6 M | 6 M | 5 M | 3 M |  |
|  | 10 | 5 M | 3 M | 3 M | 7 M | 3 M | 4 M |

F, female; M, male;

### 2.2 Radiosynthesis

[^18^F]SMBT-1 (0.74 GBq/ml) was radiosynthesized from its precursor according to a previous publication[22]. [^18^F]DPA-714 (1.48 GBq/ml) was labelled with ^18^F at its 2-fluoroethyl moiety after nucleophilic substitution of the corresponding sylate analog[25]. [^18^F]PM-PBB3 (1.48 GBq/ml) was synthesized from an automatic synthesis module and kit provided by APRINDIA therapeutics (Suzhou, China)[26]. [^18^F]florbetapir (0.56 GBq/ml) was radiosynthesized from its precursor in a fully automated procedure suitable for routine clinical application[27]. [^18^F]FDG (1.48 GBq/ml) was prepared in the radiochemistry facility of the PET Center, Huashan Hospital, Fudan University for clinical use under Good Manufacturing Practices requirements. [^11^C]PIB (0.074 GBq/ml) was radiosynthesized according to a previously described protocol[28]. The identities of the aforementioned final products were confirmed by comparison with the high-performance liquid chromatography (HPLC) retention time of the nonradioactive reference compound by coinjection using a Luna 5 μm C18(2) 100 Å (250 mm×4.6 mm) column (Phenomenex) using acetonitrile and water (60:40) solvent with a 1.0 mL/min flow rate. Radiochemical purity >95% was achieved for all aforementioned tracers.

### 2.3 MicroPET

PET experiments using [^18^F]SMBT-1, [^18^F]florbetapir, [^18^F]PM-PBB3, [^18^F]FDG, and [^18^F]DPA-714 were performed using a Siemens Inveon PET/CT system (Siemens Medical Solutions, United States)[29]. Prior to the scans, mice were anaesthetized using isoflurane (1.5%) in medical oxygen (0.3-0.5 L/min) at room temperature with an isoflurane vaporizer (Molecular Imaging Products Company, USA). The mice were positioned in a spread-supine position on the heated imaging bed and subjected to inhalation of the anaesthetic during the PET/computed tomography (CT) procedure. The temperature of the mice was monitored. A single dose of tracer (∼0.37 MBq/g body weight, 0.1–0.2 mL) was injected into the animals through the tail vein under isoflurane anaesthesia. For dynamic PET, the raw PET data were binned into 9 frames (9×600 s). Static PET/CT imaging was obtained for 10 min post intravenous administration, depending on the tracer [^18^F]FDG at 60 min, [^18^F]florbetapir at 50 min, [^18^F]SMBT-1 at 60 min, [^18^F]DPA-714 at 40 min, and [^18^F]PM-PBB3 at 90 min. PET/CT images were reconstructed using the ordered subsets expectation maximization 3D algorithm (OSEM3D), with a matrix size of 128×128×159 and a voxel size of 0.815 mm×0.815 mm×0.796 mm. Data were reviewed using Inveon Research Workplace software (Siemens). Attenuation corrections derived from hybrid CT data were applied.

### 2.4 PET data analysis

Images were processed and analysed using PMOD 4.4 software (PMOD Technologies Ltd., Switzerland). Radioactivity is presented as the standardized uptake value (SUV) (decay-corrected radioactivity per cm^3^ divided by the injected dose per gram body weight). The time−activity curves were deduced from specific volumes of interest that were defined based on a mouse MRI T_2_-weighted image template[82]. Brain regional SUVR was calculated using the cerebellum (CB) as the reference region. The mask was applied for signals outside the brain volumes of interest for illustration.

### 2.5 Immunohistochemistry

Mice were perfused under ketamine/xylazine/acepromazine maleate anaesthesia (75/10/2 mg/kg body weight, i.p. bolus injection) with ice-cold 0.1 M phosphate buffered saline (PBS, pH 7.4) and in 4% paraformaldehyde (PFA) in 0.1 M PBS (pH 7.4), fixed for 24 h in 4% PFA and then stored in 0.1 M PBS at 4°C. Coronal brain sections (40 mm) were cut around bregma 0 to -2 mm. Sections were first washed in PBS 3×10 min, followed by antigen retrieval for 20 min in citrate buffer at room temperature. Then, the sections were permeabilized and blocked in 5% normal donkey or goat serum and 1% Triton-PBS for one hour at room temperature. Free-floating tissue sections were incubated with primary antibodies against 6E10, complement component C3d (C3D), CD68, AT-8, GFAP, glucose transporter type-1 (Glut1), MAO-B and TSPO overnight at 4°C (**Suppl. Table 1**) [30, 31] and with respective secondary antibodies. Sections were incubated for 15 min in 4’,6- diamidino-2-phenylindole (DAPI), washed 2×10 min with PBS, and mounted with VECTASHIELD vibrance antifade mounting media (Vector Laboratories, Z J0215). The brain sections were imaged at ×20 magnification using an Axio Oberver Z1 slide scanner (Zeiss, Germany) using the same acquisition setting for all slices and at ×10 and ×63 magnification using a Leica SP8 confocal microscope (Leica, Germany). The images were analysed by a person blinded to the genotype using Qupath and ImageJ (NIH, U.S.A.).

### 2.6 Statistics

Two-way ANOVA with Sidak post hoc analysis was used for comparisons between groups (GraphPad Prism 9.0, CA, USA). Pearson correlation analysis was used for the analysis of the association between the regional SUVR of different tracers.

## 3 Results

### 3.1 Increased [^18^F]SMBT-1 SUVR in the cortex and hippocampus of 10-month-old APP/PS1 mice

The HPLC and quality control (QC) of [^18^F]SMBT-1 are shown in **SFig. 1**. The time activity curve of [^18^F]SMBT-1 indicated that 60 min postinjection in mice is suitable for static scans, which is similar to the application in the clinical setting (**SFig. 2**). We observed that the [^18^F]SMBT-1 SUVR (Cb as reference region) of MAO-B, indicative of reactive astrocytes, was higher in the cortex and hippocampus of 10-month-old APP/PS1 mice than in 5-month-old APP/PS1 and age-matched wild-type mice. No regional difference in [^18^F]SMBT-1 SUVR was observed between 11-month-old 3×Tg mice and age-matched wild-type mice (**Figs. 1a-f**).

**Fig. 1.**
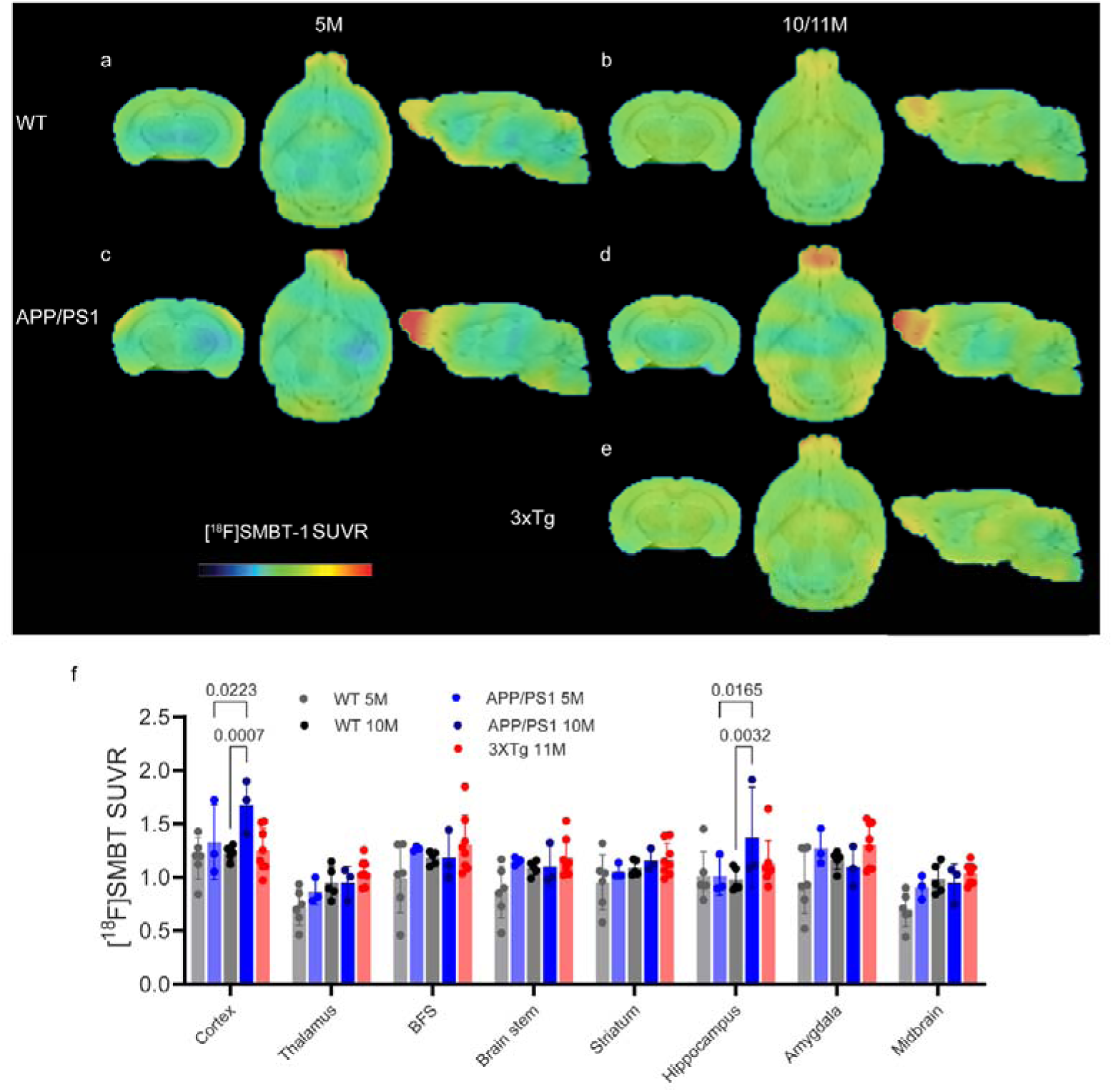
Increased [^18^F]SMBT-1 brain uptake in 10-month-old APP/PS1 mice compared to age-matched wild-type mice. (**a-e**) SUVR images of 5- and 10-month-old WT (a, b), 5- and 10-month-old APP/PS1 (c, d), and 11-month-old 3×Tg mice (e). SUVR scale 0-2.2. (**f**) Quantification of [^18^F]SMBT-1 using the cerebellum as a reference region in WT, APP/PS1 and 3×Tg mice. BFS, basal forebrain system;

### 3.2 Higher [^18^F]florbetapir SUVR in the cortex and hippocampus of APP/PS1 mice from 5 months of age

The [^18^F]florbetapir SUVR (Cb as reference region) was higher in the thalamus, basal forebrain system, brain stem and midbrain of 5-month-old APP/PS1 mice than in age-matched wild-type mice. Higher [^18^F]florbetapir SUVR was observed in the cortex and hippocampus in 10-month-old APP/PS1 mice than in age-matched wild-type mice and 5-month-old APP/PS1 mice (**Fig. 2**). In contrast, no regional difference in either [^18^F]florbetapir or [^11^C]PIB SUVR was observed in the brains of 11-month-old 3×Tg mice compared to age-matched wild-type mice (**Figs. 2e, f**, **g**).

**Fig. 2.**
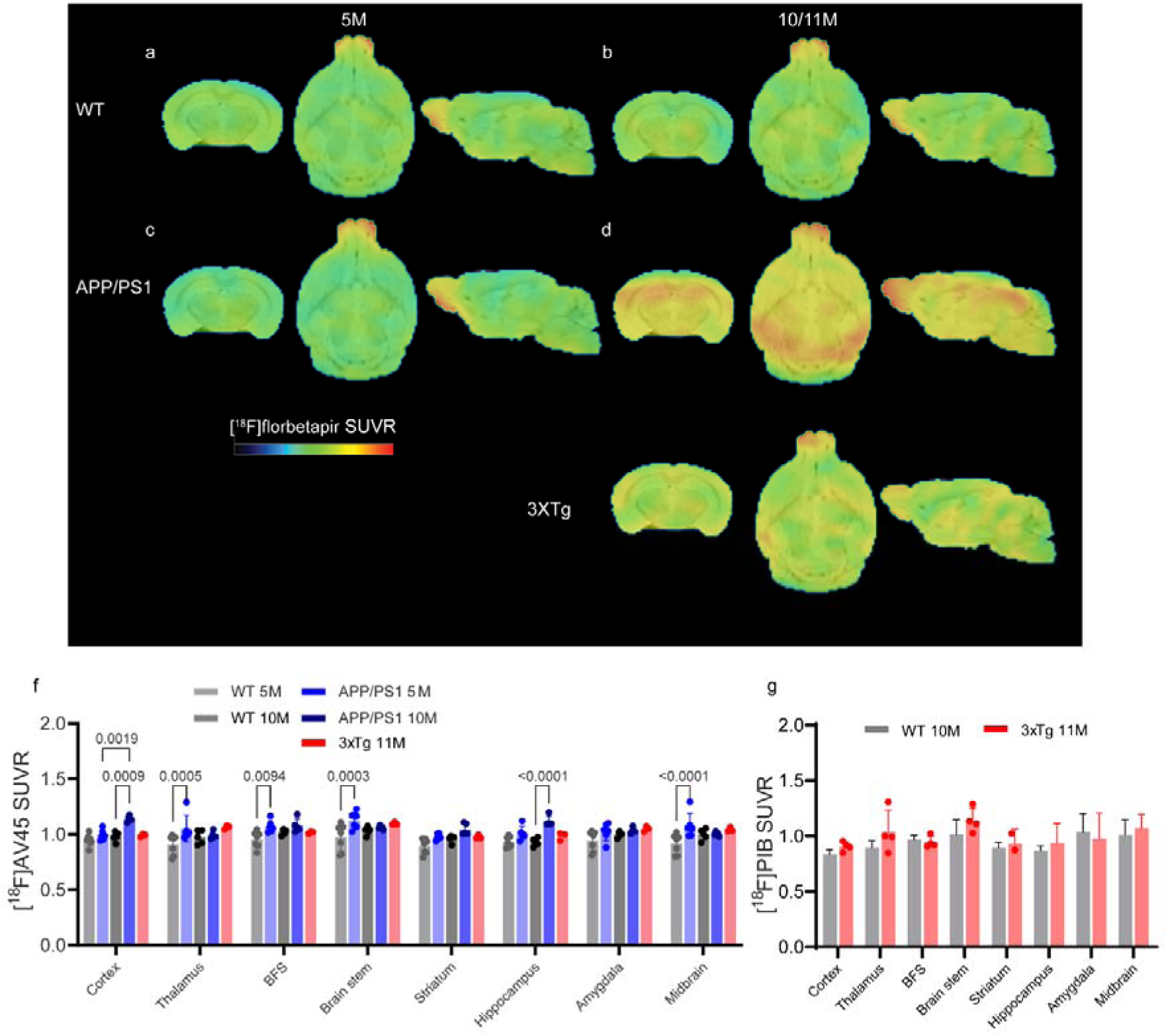
Increased [^18^F]florbetapir brain uptake in 10-month-old APP/PS1 mice compared to 5-month-old APP/PS1 mice and age-matched wild-type mice and 3×Tg mice. (**a-e**) SUVR images of 5- and 10-month-old WT (a, b), 5- and 10-month-old APP/PS1 (c, d), and 11-month-old 3×Tg mice (e). SUVR scale 0-2.2. (**f**) Quantification of [^18^F]florbetapir using the cerebellum as a reference region in WT, APP/PS1 and 3×Tg mice. (**g**) No difference in [^11^C]PIB brain uptake in the 3×Tg mice compared to wild-type mice. The SUVR was calculated using the cerebellum as the reference brain region. BFS, basal forebrain system; BFS, basal forebrain system;

### 3.3 No difference in **[^18^F]**PM-PBB3 SUVR in the cortex and hippocampus of APP/PS1 mice and 3×Tg mice

We chose 90 min postinjection for the [^18^F]PM-PBB3 static scan based on the time activity curve (**SFig. 3**) and previous studies. The cerebellum was used as the reference brain region as in previous PET studies with [^18^F]PM-PBB3[32]. No regional difference in [^18^F]PM-PBB3 SUVR was observed in the brains of 5-month-old APPPS1 mice or 11-month-old 3×Tg mice compared to age-matched wild-type mice (**Fig. 3**).

**Fig. 3.**
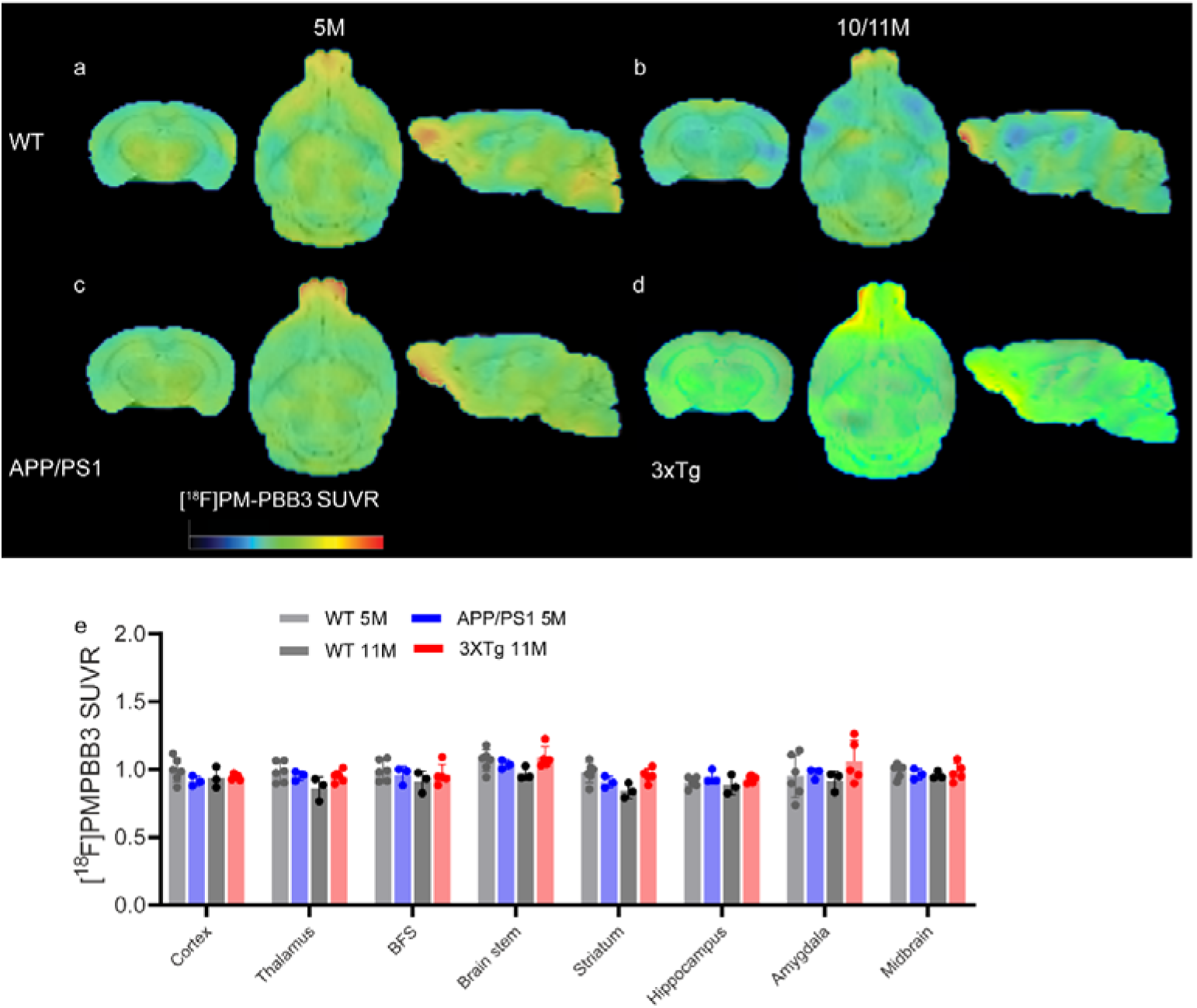
No difference in [^18^F]PM-PBB3 brain uptake in 3×Tg mice and age-matched wild-type mice. (**a-e**) SUVR images of 5- and 10-month-old WT (a, b), 5-month-old APP/PS1 (c), and 11-month-old 3×Tg mice (d). SUVR scale 0-2.2. (e) Quantification of [^18^F]PM-PBB3 using the cerebellum as a reference brain region in WT and 3×Tg mice. BFS, basal forebrain system;

### 3.4 Reduced [^18^F]FDG SUVR in the amygdala of APP/PS1 mice at 5 months of age

The [^18^F]FDG SUVR (Cb as reference region) was lower in the thalamus, basal forebrain system, striatum, and midbrain of 5-month-old APP/PS1 mice than in age-matched wild-type mice. Reduced [^18^F]FDG SUVR was observed in the amygdala of 10-month-old APP/PS1 mice compared with 5-month-old APP/PS1 mice (**Fig. 4**). No regional difference in [^18^F]FDG SUVR was observed in the brains of 11-month-old 3×Tg mice compared to age-matched wild-type mice (**Fig. 4**).

**Fig. 4.**
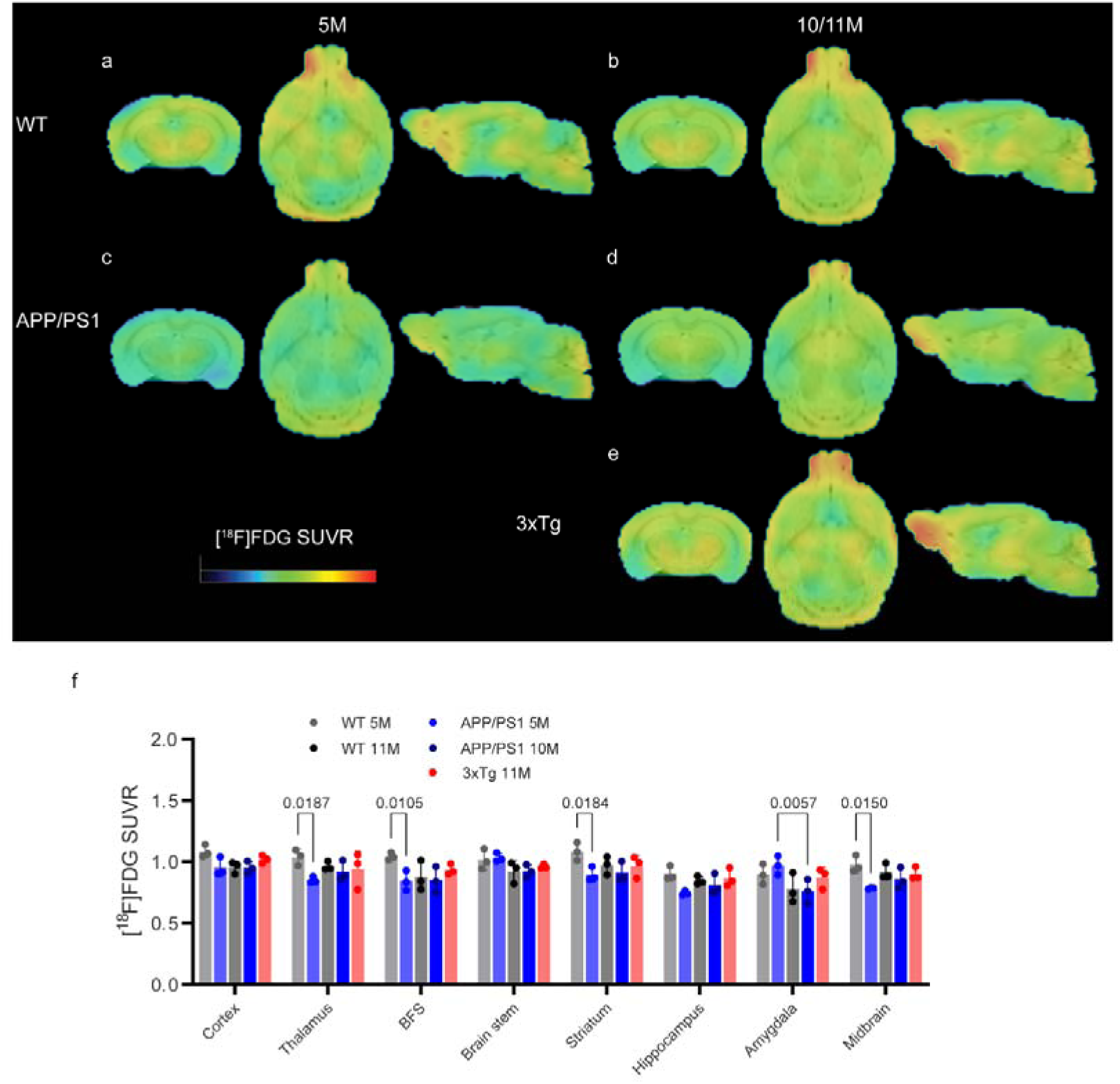
Reduced [^18^F]FDG brain uptake in 5-month-old APP/PS1 mice compared to age-matched wild-type mice. (**a-e**) SUVR images of 5- and 10-month-old WT (a, b), 5- and 10-month-old APP/PS1 (c, d), and 11- month-old 3×Tg mice (e). SUVR scale 0-1.8. (**f**) Quantification of [^18^F]FDG using the cerebellum as a reference brain region in WT, APP/PS1 and 3×Tg mice. Correlation between [^18^F]SMBT-1 SUVR and [^18^F]DPA-714 SUVR in the cortex. BFS, basal forebrain system;

### 3.5 [^18^F]DPA-714 SUVR did not differ between APP/PS1 mice, 3×Tg mice and wild-type mice

We used the cerebellum as a reference brain region for [^18^F]DPA-714 according to earlier studies using [^18^F]DPA-714 and [^11^C]PBR28 in mouse models [33]. No difference in [^18^F]DPA-714 SUVR was observed in 5- or 10-month-old APP/PS1 mice compared to age-matched wild-type mice (**Fig. 5**). No difference in [^18^F]DPA-714 SUVR was observed in 11-month-old 3×Tg mice compared to age-matched wild-type mice.

**Fig. 5.**
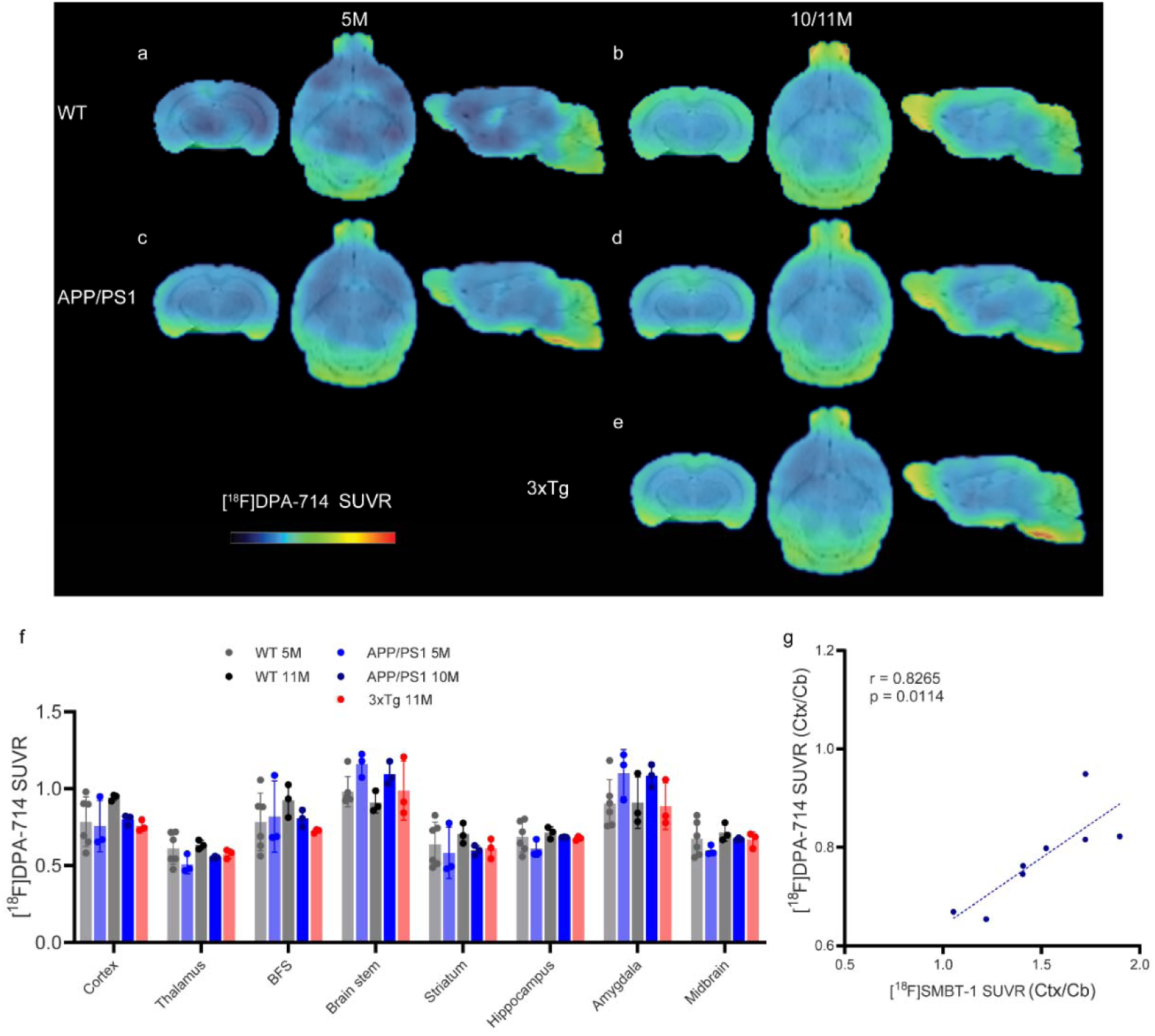
Comparable [^18^F]DPA-714 brain uptake in 5-month- and 10-month-old APP/PS1 mice, age- matched wild-type mice and 3×Tg mice. (**a-e**) SUVR images of 5- and 10-month-old WT (a, b), 5- and 10- month-old APP/PS1 (c, d), and 11-month-old 3×Tg mice (e). SUVR scale 0-2.2. (**f**) Quantification of [^18^F]DPA- 714 using the cerebellum as a reference brain region in WT, APP/PS1 and 3×Tg mice. BFS, basal forebrain system;

### 3.6 Regional correlation between [^18^F]SMBT-1, [^18^F]florbetapir, [^18^F]PM-PBB3, [^18^F]FDG, and [18F]DPA-714

To assess the spatial association between different pathologies, Pearson’s correlation analysis was performed for [^18^F]SMBT-1, [^18^F]florbetapir [^18^F]PM-PBB3, [^18^F]FDG and [^18^F]DPA-714. A positive correlation was observed between the cortical [^18^F]SMBT-1 SUVR and [^18^F]DPA-714 SUVR in the cortex but not in the hippocampus (**Fig. 5g**).

### 3.7 Reactive astrocytes and microgliosis with tau inclusions and A**β** deposits

Next, we evaluated the distribution of MAO-B and TSPO along with the astrocytic markers GFAP/C3d and microglial markers CD68, phospho-tau (AT-8), Aβ deposits (6E10), and GluT1 in brain tissue slices (**Figs. 6-8, SFig. 4**). MAO-B expression was detected on astrocytes with colonization of C3D immunofluorescence, which was highly upregulated in A1 reactive astrocytes in the cortex and hippocampus of APP/PS1 and 3×Tg mice (**Figs. 5b, c,** zoom-in). Colocalization of TSPO with CD68 (phagocytic microglia) and GFAP-positive astrocytes was observed in the cortex and hippocampus (**Figs. 6b, c**). In addition, colocalization of TSPO with AT-8-positive phospho-tau inclusions was observed in the hippocampus of 3×Tg mice (**Fig. 6d**). In 3×Tg mice, sporadic Aβ deposits (mainly intracellular) and tau inclusions were observed, mainly in the subiculum and hippocampus (**Fig. 8**), validating the lack of amyloid and tau PET updates in this strain. The level of the glucose transport protein GluT1 appeared comparable in the cortex and hippocampus from 3×Tg mice and wild-type mice (**SFig. 4**).

**Fig. 6.**
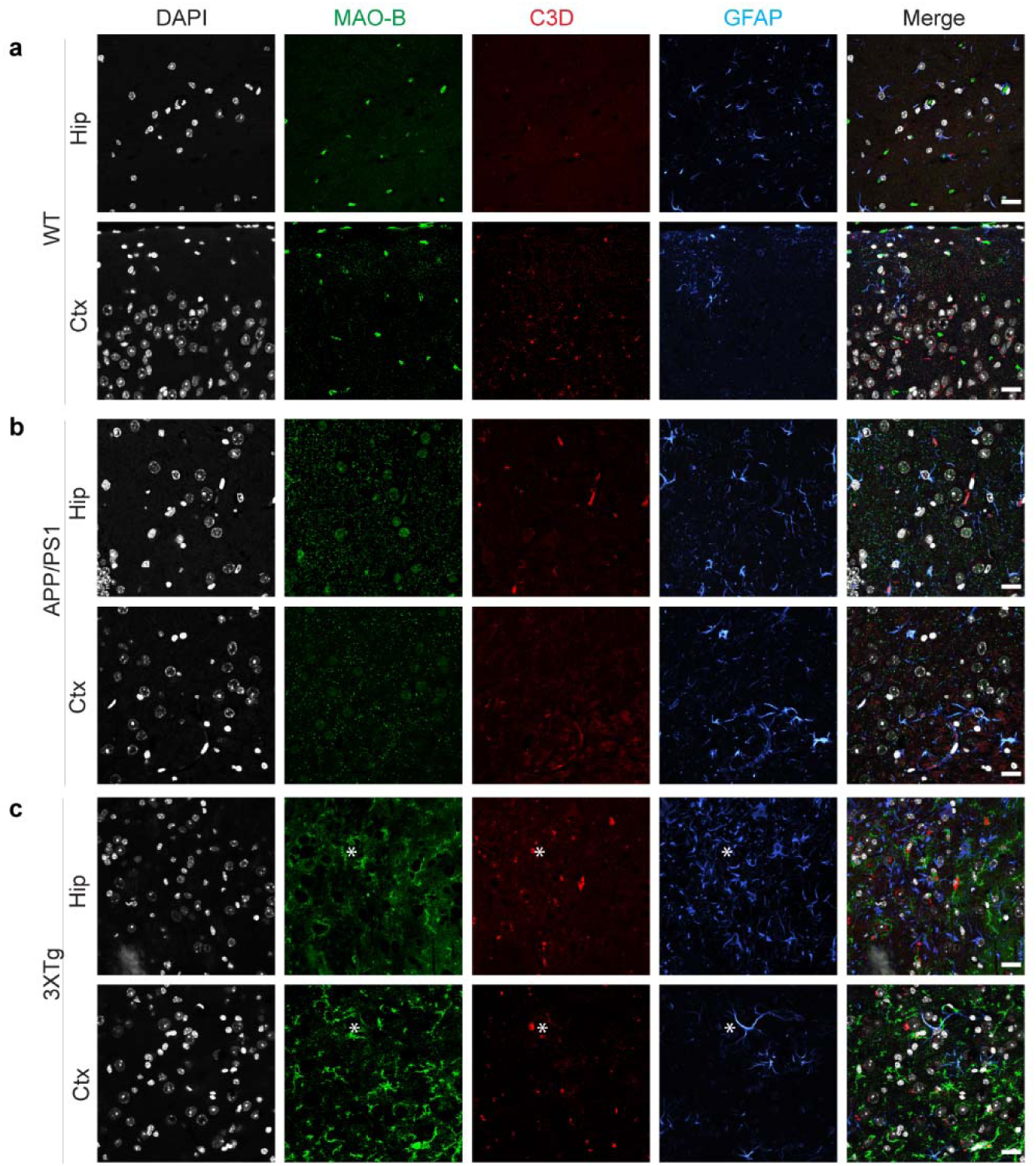
Immunofluorescence staining of MAO-B with astrocyte markers in the cortex and hippocampus of the mouse brain. (**a-c**) Brain tissue sections of WT, APP/PS1 and 3×Tg mice were stained for MAO-B (green)/C3D (red)/GFAP (blue) and showed colocalization of MAO-B on C3D-positive astrocytes in the Ctx and Hip. Nuclei were counterstained with DAPI (gray). * indicates colocalization. Scale bar=20 μm.

**Fig. 7.**
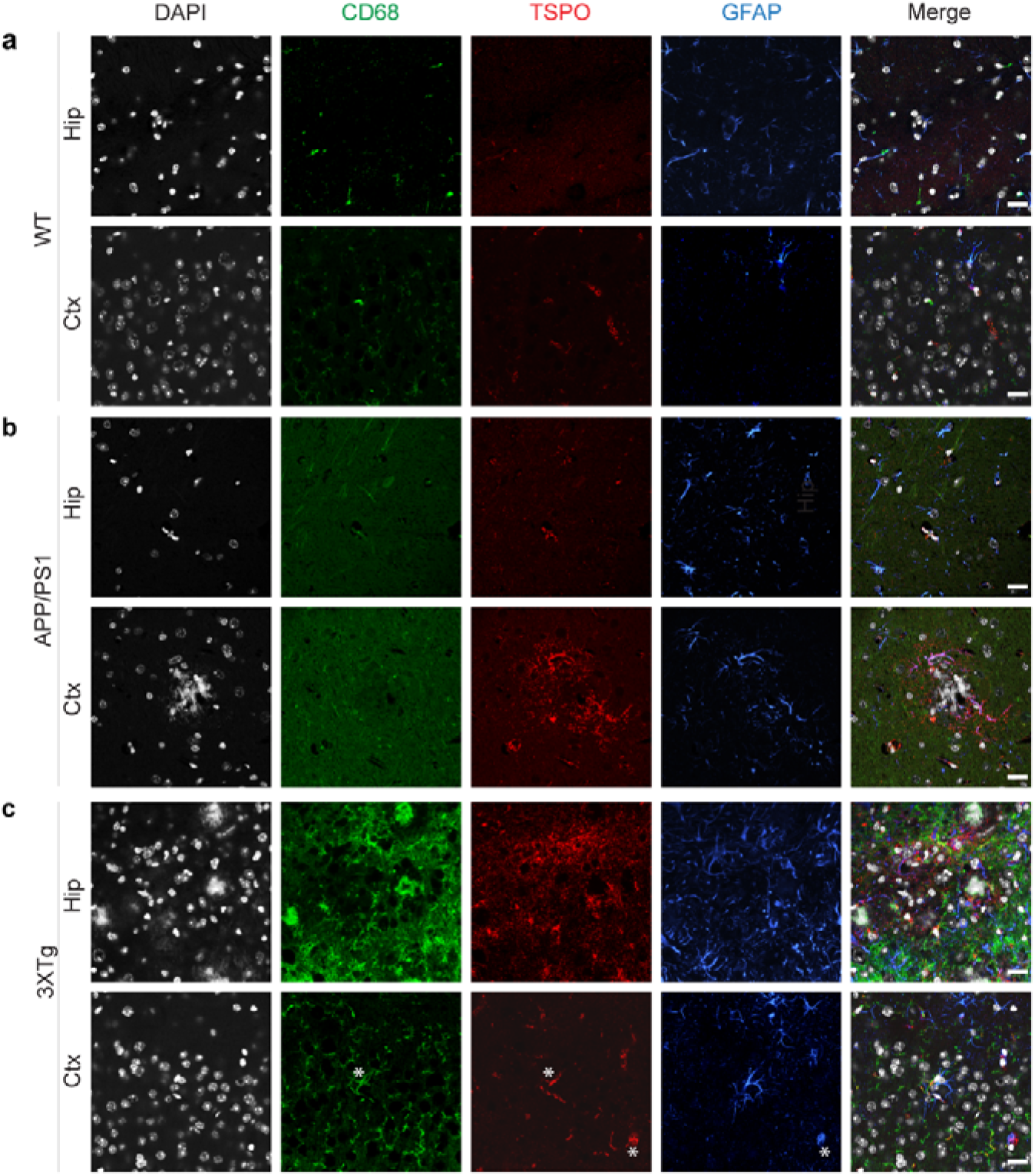
Immunofluorescence staining of TSPO with astrocyte and microglial markers in the cortex and hippocampus of the mouse brain. (**a-c**) Brain tissue sections of WT, APP/PS1 and 3×Tg mice were stained for CD68 (green)/TSPO (red)/GFAP (blue) to show the localization of TSPO on astrocytes and microglia in the Ctx and Hip. Nuclei were counterstained with DAPI (gray). *indicates colocalization. Scale bar=20 μm.

**Fig. 8.**
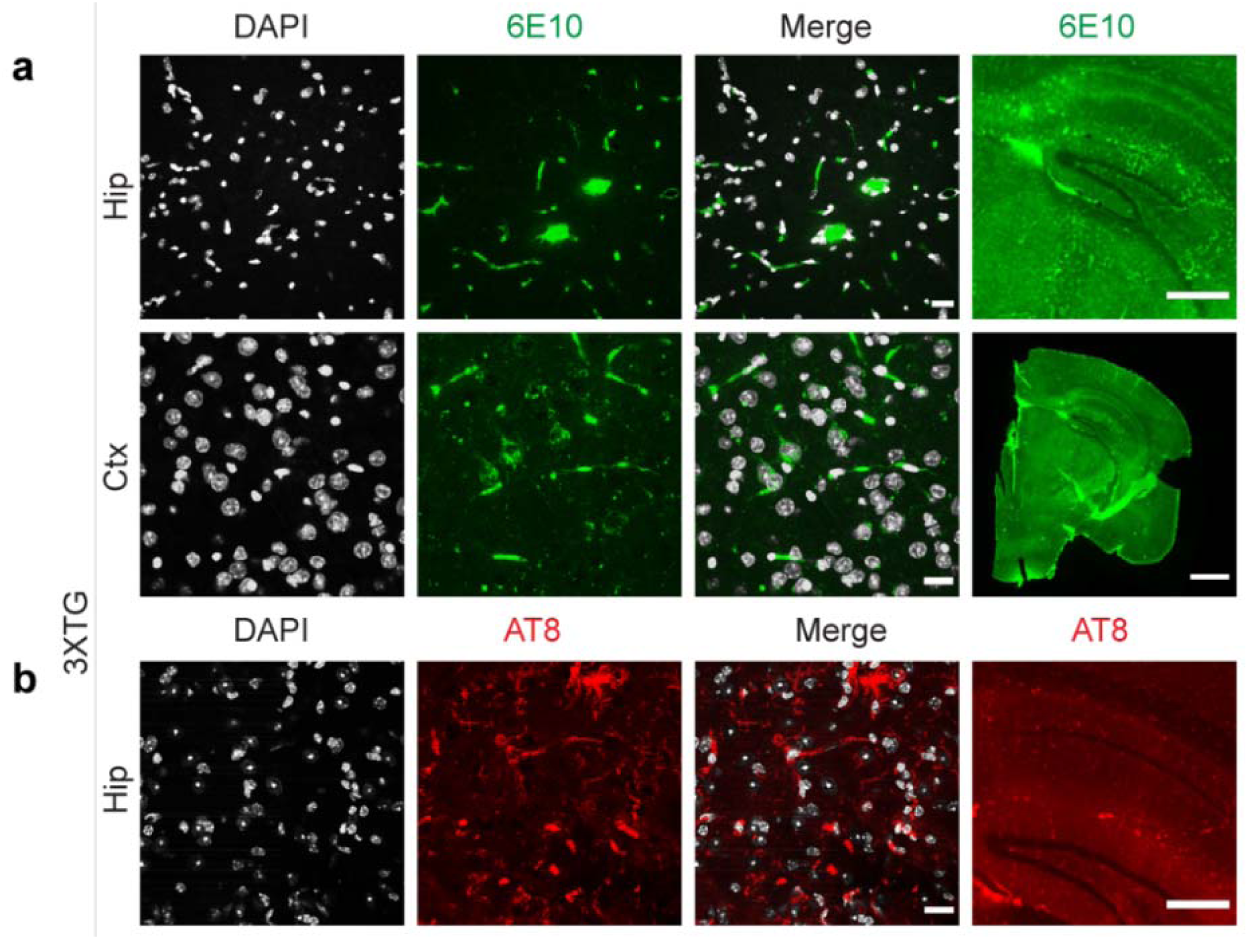
Limited amyloid-beta plaque and tau deposits in the brains of 3×Tg mice. (**a-c**) Limited amyloid and tau inclusion was validated in the subiculum and cortex brain tissue sections of 3×Tg mice stained for 6E10 (mainly intracellular, green, a) and AT-8 (red, b) in the hippocampus. Nuclei were counterstained with DAPI (white). Scale bar=1 mm (column 4 in the second row of a) and 500 μm (column 4 in the first row of a and b), 20 μm for the others.

## Discussion

Here, we demonstrated increased brain regional [^18^F]SMBT-1 and [^18^F]florbetapir brain uptake in 10-month-old APP/PS1 mice and reduced regional [^18^F]FDG and comparable [^18^F]DPA-714 SUVR in 5-month-old APP/PS1 mice compared to age-matched WT mice. Moreover, [^18^F]SMBT-1 and [^18^F]DPA-714 uptake correlated only in the cortex, showing divergent regional patterns of reactive astrocytes and microgliosis.

Here, we found increased cortical and hippocampal MAO-B levels by [^18^F]SMBT-1 PET in 10-month-old APP/PS1 mice compared to wild-type mice, which is supported by immunohistochemical staining results. Our [^18^F]SMBT-1 imaging results are in line with earlier reports using MAO-B tracers in APP models: higher [^18^F]F-DED uptake has been reported in the thalamus of PS2APP mice at 5, 13, and 19 months and in the hippocampus at 14 and 19 months compared to wild-type mice [11]. [^11^C]DED measured increased uptake in 6- month-old APPswe mice preceding amyloid plaque deposition using [^11^C]AZD2184 [10].

For amyloid imaging, PET studies using [^18^F]florbetapir [38–41], [^18^F]florbetaben, [^11^C]PIB [42, 43], and [^18^F]fluotemetamol in APP/PS1 mice have been reported. Our finding of increased [^18^F]florbetapir SUVR in the cortex and hippocampus was in line with known Aβ aggregate distribution and immunofluorescence staining in the brains of APP/PS1 mice. For tau imaging, PET using [^11^C]PBB3, [^18^F]PM-PBB3, and [^18^F]PI-2620 in PS19 and rTg4510 mice [31, 32, 45–51], as well as [^11^C]THK5317 [52] and [^11^C]THK5117 [53], which bind to both tau and MAO-B, in double mutant TgF344 rats has been reported. We observed no change in the [^18^F]PM-PBB3 SUVR in the hippocampus and cortex of APP/PS1 mice at 5 months. Thus far, only one [^18^F]flortaucipir *ex vivo* autoradiography study has been reported in aged APP/PS1 mouse brain slices [54].

Inconsistent results have been reported for [^18^F]FDG updates in animal models of AD, partly due to the difference in the imaging protocol, aging and heterogeneity between animals [55]. Here, we found that there was a reduced level of [^18^F]FDG in 5-month-old APP/PS1 mice compared to wild-type mice, which was absent at 10 months. A longitudinal study in APP/PS1-21 mice showed reduced cerebral glucose metabolism using [^18^F]FDG and increased [^18^F]DPA-714 from 12 months [64].

For microglial imaging, tracers targeting TSPO [56, 57] have been most widely used, including 1^st^ generation [^11^C]PK11195, 2^nd^ generation [^18^F]DPA-714, [^11^C]PBR28, and 3^rd^ generation [^18^F]GE-180. [^18^F]DPA-714 showed favorable binding potential and selectivity and low nonspecific binding compared to [^11^C]PK11195 [58]. We observed no difference in the brain regional [^18^F]DPA-714 SUVR (Cb as reference region) in 5- or 10- month-old APP/PS1 mice compared to wild-type mice. In line with our findings, several imaging studies using the three generations of TSPO tracers have been reported in APP/PS1 mice only at a later age [59]: increased [^11^C]PK11195 uptake at 16-19 months (not at 13-16 months) [60]; higher [^18^F]GE180 at 26 months compared to 4 months [61]; higher [^18^F]DPA-714 uptake using Cb as the reference region at 12 months [63]/18 months [62], and at 12-16 months using muscle as the reference region [25].

We found no difference in the levels of [^18^F]SMBT-1(MAO-B), [^18^F]florbetapir/[^11^C]PIB(Aβ), [^18^F]PM- PBB3(tau), [^18^F]FDG and [^18^F]DPA-714(TSPO) SUVRs in 3**×**Tg mice compared to aged-matched controls at 11 months. Our results are in line with a recent study showing no difference using [^11^C]DED in the hippocampus and cortex of 10-month-old 3**×**Tg mice, although an increase in [^18^F]sulforhodamine-101 [14](different target) was observed. Our observation of a lack of increase in amyloid and TSPO uptake is in line with previous observations using [^11^C]PIB([^18^F]florbetaben) [65] and [^11^C]PK11195 in 3**×**Tg mice at 4-16 months [66, 67]. However, another study with TSPO and amyloid SPECT imaging showed an increase in the hippocampus in 3**×**Tg mice at 10 months compared to wild-type mice [68]. We observed that there were no redistribution or intensity changes in GFAP, MAO-B, C3d, CD68, TSPO, or GluT1 and a limited distribution of Aβ and tau immunoreactivity in 11-month-old 3**×**Tg mice, which is in line with recent systematic characterization studies in 3**×**Tg mice [69, 70].

Reactive astrocytes acquire neuroprotective as well as deleterious signatures in response to tau and Aβ pathology [71] and influence the effects of amyloid-β on tau pathology in preclinical AD [72]. The [^11^C]BU99008 measure of astrocyte reactivity correlated with a reduction in cerebral glucose metabolism, gray matter volume and amyloid load in cognitively impaired individuals [75]. Here, we observed an increased level of [^18^F]SMBT-1 in MAO-B along with an increase in Aβ and tau accumulation in the hippocampus and cortex of APP/PS1 mice, preceding microgliosis measured by [^18^F]DPA-714. Cortical [^18^F]SMBT-1 SUVR correlated positively with [^18^F]DPA-714 SUVR but not in other brain regions. The association between neurotoxic reactive astrocytes is induced by activated microglia in mouse models [73]. In APP/PS1 mice, upregulated levels of MAO-B and reactive astrocytes have been shown to increase the level of tau inclusions, neuronal death, brain atrophy, and impaired spatial memory ability in a H_2_O_2_-dependent manner [37]. In addition, MAO-B reversibly increased astrocytic γ-aminobutyric acid production in reactive astrocytes [35], which was associated with synaptic and memory impairments in APP/PS1 mice [36]. Previous topological analyses revealed that astrocytes respond to plaque-induced neuropil injury primarily by changing phenotype and hence function rather than location [34].

There are several limitations in this study. 1) The mice examined were cross-sectional, not longitudinal. 2) The sample size and sex balance of the animals were not optimal. 3) We did not provide detailed profiling of the heterogeneity of astrocytes and microglia, such as by using transcriptomics. Notably, there are distinct dynamic profiles of microglial activation [76] and reactive astrocytes between human and mouse models [77]. Therefore, further detailed profiling and analysis are needed.

## Conclusion

Here, we showed increased levels of [^18^F]SMBT-1 and [^18^F]florbetapir in the brains of 10-month-old APP/PS1 mice compared to age-matched wild-type mice, preceding changes in the level of [^18^F]DPA-714. Given that reactive astrocytes are emerging treatment targets for AD [78], small-animal reactive astrocyte PET can provide in vivo tools for outcome measures in these therapeutic developments [39].

## Declaration

### Author contributions

The study was designed by KY and RN. KY performed radiosynthesis, HPLC and microPET. CM performed staining, microscopy and miceoscopy image analysis. RN performed the microPET analysis. YK, CM, and RN wrote the first draft. All authors contributed to the revision of the manuscript. All the authors have read and approved the final manuscript.

### Data availability

The datasets generated and/or analysed during the current study are available from the corresponding author upon reasonable request.

### Ethics approval

The PET imaging and experimental protocol was approved by the Institutional Animal Care and Ethics Committee of Fudan University and performed in accordance with the National Research Council’s Guide for the Care and Use of Laboratory Animals. All experiments in Zurich were performed in accordance with the Swiss Federal Act on Animal Protection and were approved by the Cantonal Veterinary Office Zurich (ZH162/20).

### Consent to participate

Not applicable.

### Consent to publish

Not applicable.

---

**Supplementary Fig 1.**
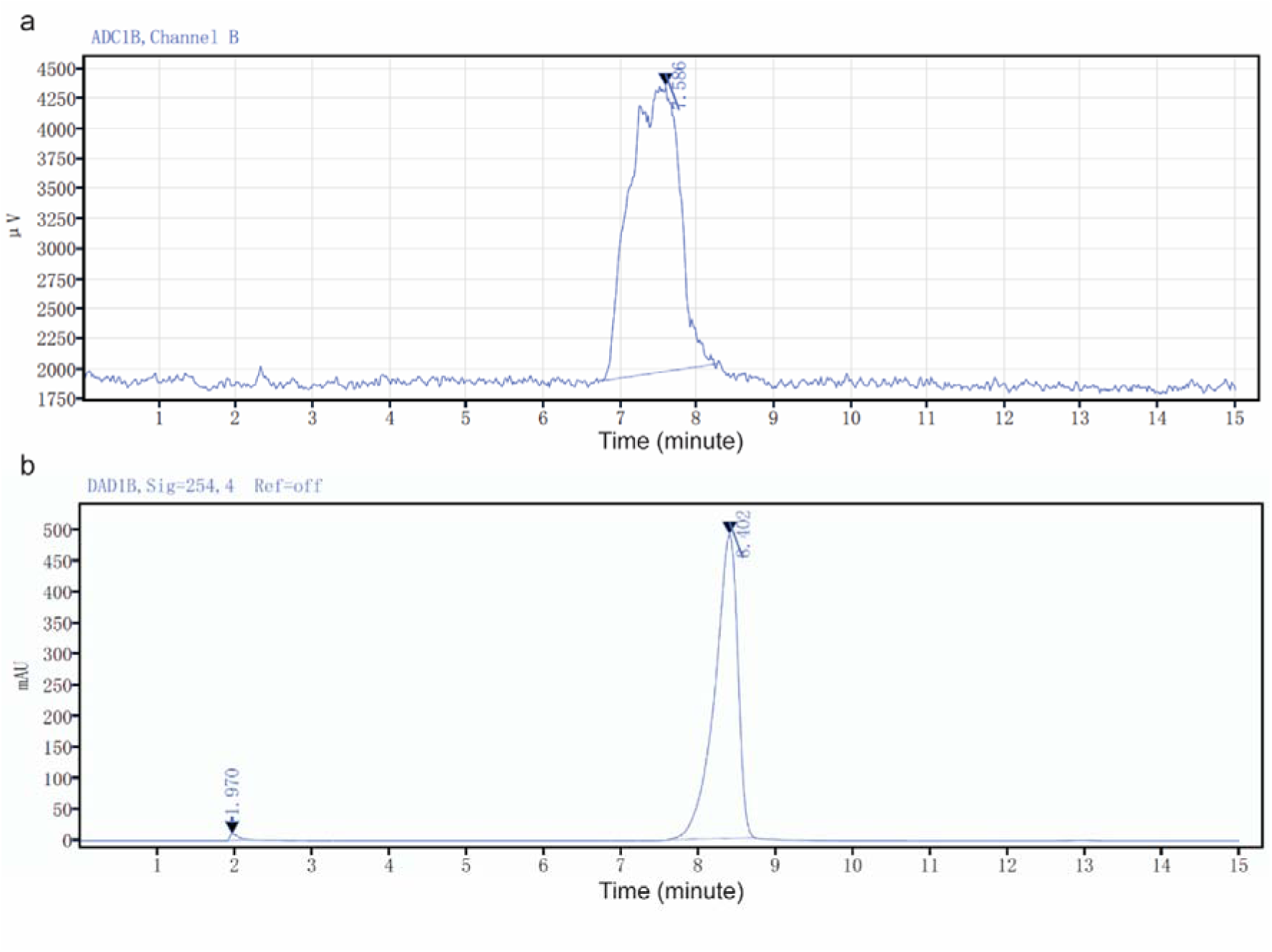
HPLC chromatogram of synthesized [^18^F]SMBT-1 and standard. **(a)** Synthesized SMBT-1, (**b**) Standard.

**Supplementary Fig 2.**
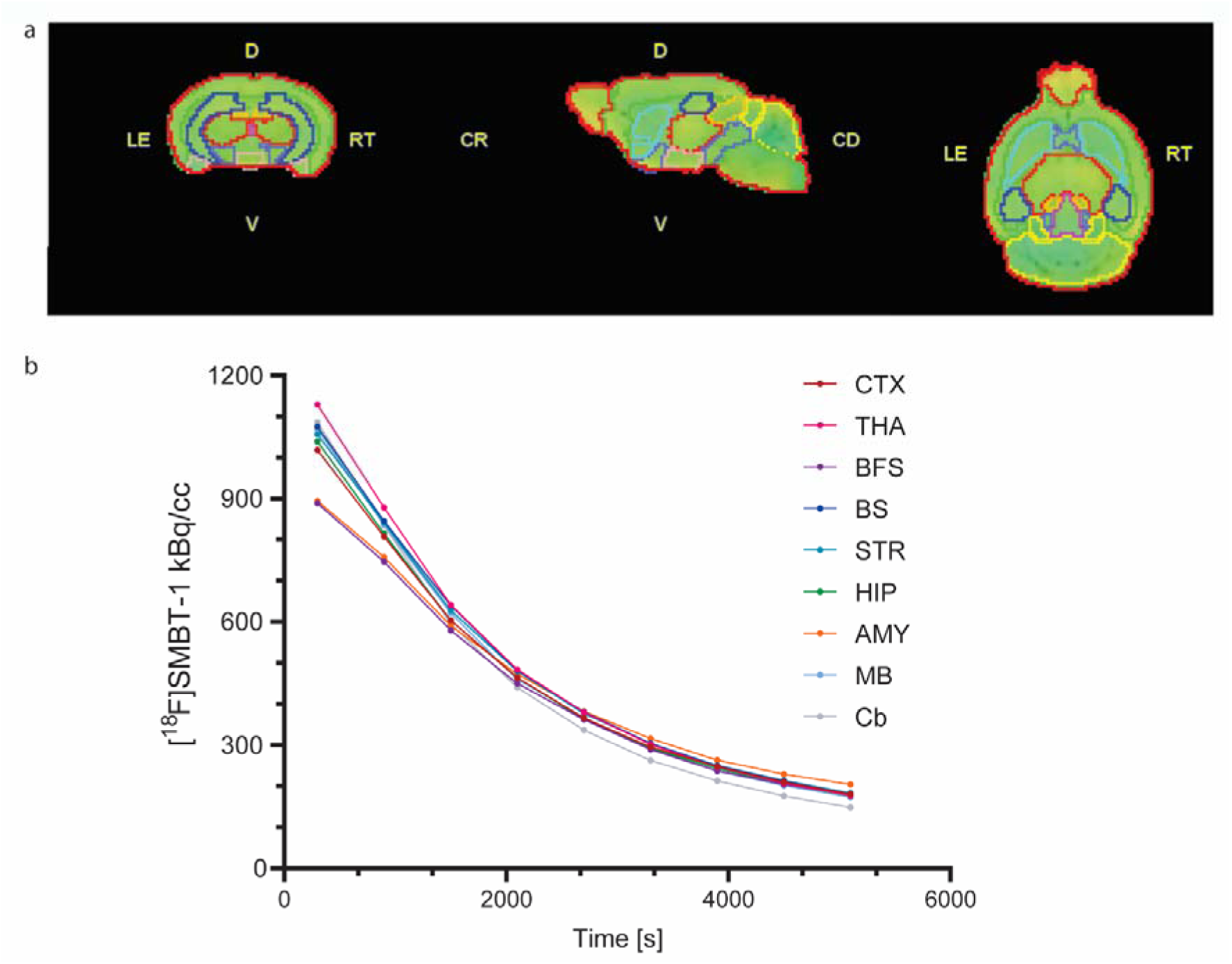
Volume-of-interest analysis and time activity curve of [^18^F]SMBT-1 in a mouse brain. (**a**) Coronal, sagittal and horizontal views of a mouse brain. (**b**) Regional [^18^F]SMBT-1 time activity (kBq/cc) in the brain. CTX, cortex; THA, thalamus; BFS, basal forebrain system; BS, brain stem; STR, striatum; HIP, hippocampus; AMY, amygdala; MB, midbrain; Cb, cerebellum.

**Supplementary Fig 3.**
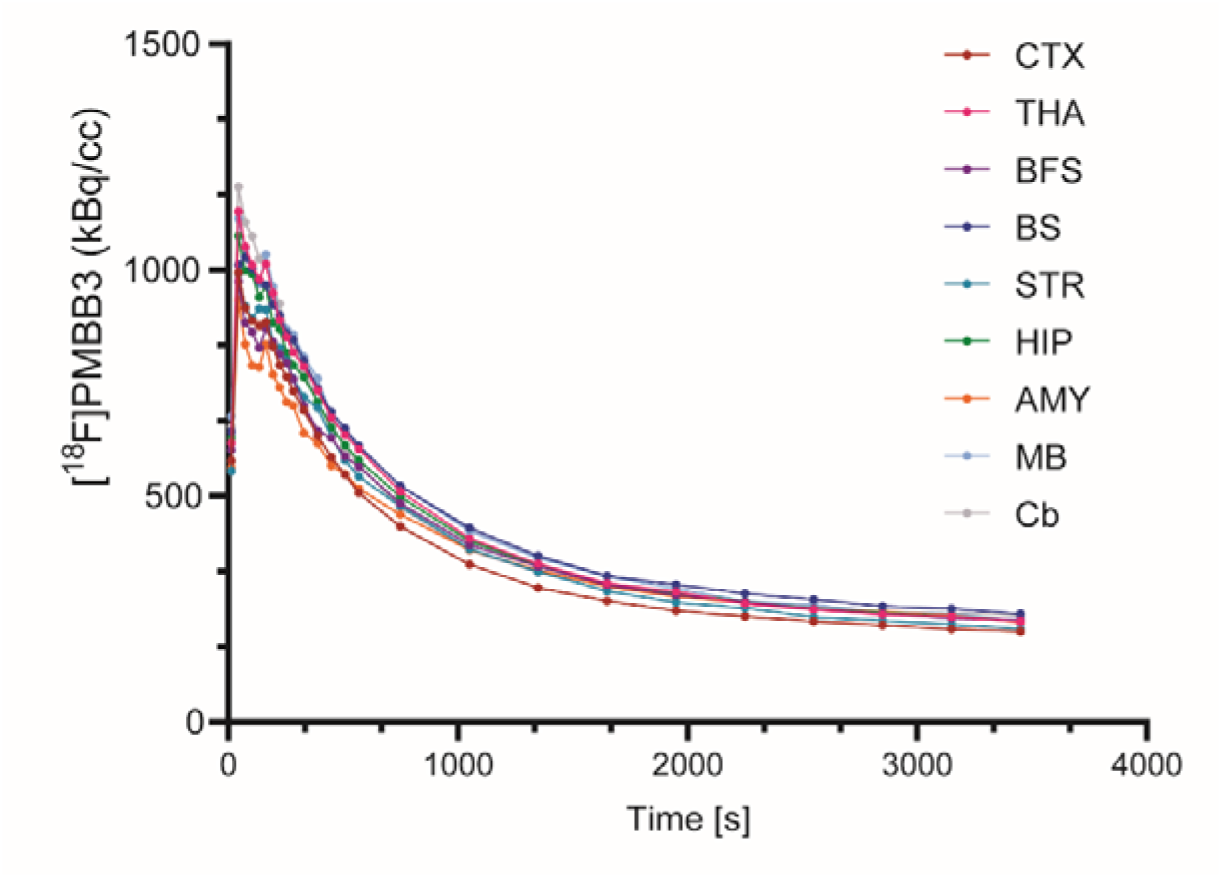
Time activity curve of [^18^F]PM-PBB3 in the mouse brain. Regional time activity (kBq/cc) in the brain CTX, cortex; THA, thalamus; BFS, basal forebrain system; BS, brain stem; STR, striatum; HIP, hippocampus; AMY, amygdala; MB, midbrain; Cb, cerebellum.

**Supplementary Fig 4.**
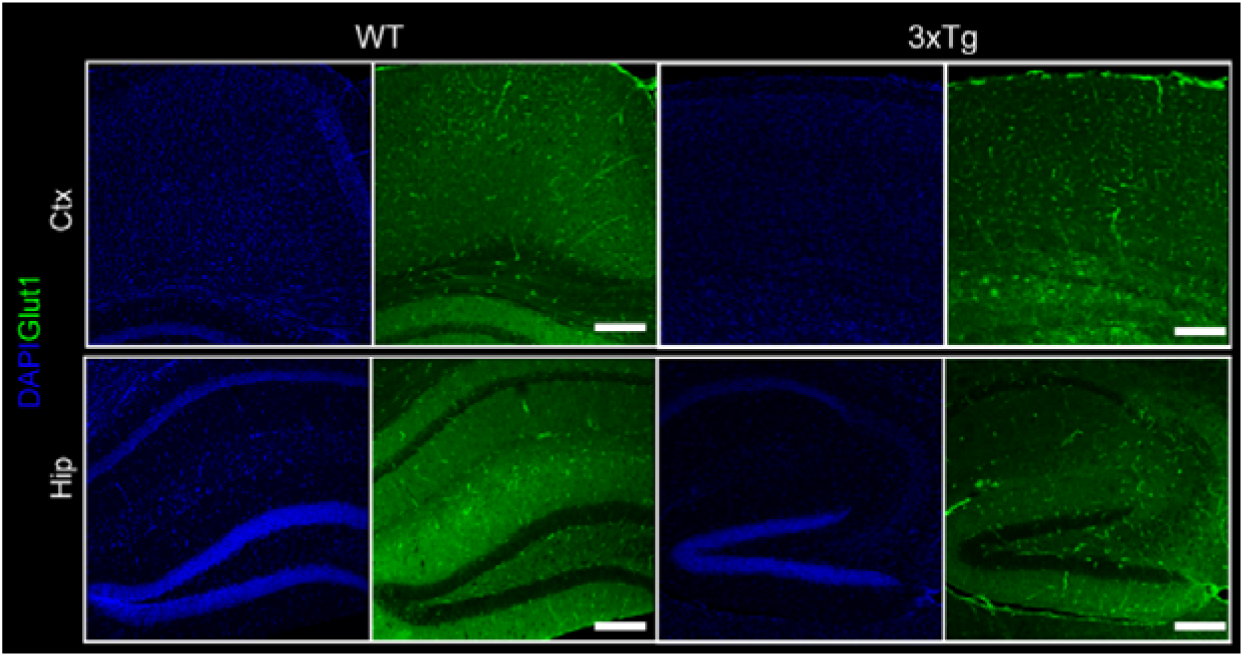
GluT1 staining in WT and 3×Tg mouse brains. (**d**) Brain tissue sections of wild-type and 3×Tg mice were stained for GluT1 (green) in the cortex and hippocampus. Nuclei were counterstained with DAPI (blue). Scale bar = 200 μm.

**Supplementary Table 1.** List of antibodies.

| Antibody | Catalog no | Dilution | Supplier |
| --- | --- | --- | --- |
| Alexa fluor488 donkey anti-mouse IgG (H+L) | 715-545-151 | 1:500/1:250/1:125 | Jackson |
| Goat anti-mouse IgG (H+L) highly cross-adsorbed secondary antibody, Alexa fluor plus 488 | A32723 | 1:500 | Invitrogen |
| Cy3 donkey anti-rabbit IgG (H+L) | 711-165-152 | 1:250 | Jackson |
| Donkey anti-goat IgG (H+L) cross-adsorbed secondary antibody, Alexa fluor 546 | A11056 | 1:200/1:50 | Invitrogen |
| Alexa fluor647 donkey anti-guinea pig IgG (H+L) | 706-605-148 | 1:250 | Jackson |
| Donkey anti-rabbit IgG (H+L) highly cross-adsorbed secondary antibody, Alexa Fluor Plus 647 | A32795 | 1:200 | Invitrogen |
| DAPI (4',6-Diamidino-2-Phenylindole, Dihydrochloride) | D1306 | 1:1000 | Invitrogen |
| Mouse phosphor-Tau (Ser202, Thr205) monoclonal antibody (AT8) | MN1020 | 1:1000 | Invitrogen |
| Mouse Purified anti-β-Amyloid, 1-16 monoclonal antibody (6E10) | 803001 | 1:1000 | Biolegend |
| Goat complement component C3d polyclonal antibody | AF2655 | 1:200 | R&D Systems |
| Guinea pig GFAP polyclonal antibody | BP5082 | 1:1000 | OriGene |
| Mouse Glut1 (C-terminus) monoclonal antibody | MABS132 | 1:1000 | Milipore |
| Rabbit PBR monoclonal antibody [EPR5384] (TSPO) | ab109497 | 1:1000 | Abcam |
| Mouse MAO-B (D-6) monoclonal antibody | sc-515354 | 1:500 | Santa Cruz |

## References

1. Verkhratsky A, Nedergaard M. Physiology of Astroglia. Physiol Rev. 2018;98:239–389.

2. Escartin C, Galea E, Lakatos A, O’Callaghan JP, Petzold GC, Serrano-Pozo A, et al. Reactive astrocyte nomenclature, definitions, and future directions. Nat Neurosci. 2021;24:312–25.

3. Heneka MT, Carson MJ, El Khoury J, Landreth GE, Brosseron F, Feinstein DL, et al. Neuroinflammation in Alzheimer’s disease. Lancet Neurol. 2015;14:388–405.

4. Beyer L, Stocker H, Rujescu D, Holleczek B, Stockmann J, Nabers A, et al. Amyloid-beta misfolding and GFAP predict risk of clinical Alzheimer’s disease diagnosis within 17 years. Alzheimers Dement. 2022.

5. Ferrari-Souza JP, Ferreira PCL, Bellaver B, Tissot C, Wang YT, Leffa DT, et al. Astrocyte biomarker signatures of amyloid-β and tau pathologies in Alzheimer’s disease. Mol Psychiatry. 2022;27:4781–9.

6. Ni R, Röjdner J, Voytenko L, Dyrks T, Thiele A, Marutle A, et al. In vitro Characterization of the Regional Binding Distribution of Amyloid PET Tracer Florbetaben and the Glia Tracers Deprenyl and PK11195 in Autopsy Alzheimer’s Brain Tissue. J Alzheimer’s Dis. 2021;80:1723–37.

7. Marutle A, Gillberg P-G, Bergfors A, Yu W, Ni R, Nennesmo I, et al. 3 H-Deprenyl and 3 H-PIB autoradiography show different laminar distributions of astroglia and fibrillar β-amyloid in Alzheimer brain. J Neuroinflammation. 2013;10:1–15.

8. Serrano-Pozo A, Mielke ML, Gómez-Isla T, Betensky RA, Growdon JH, Frosch MP, et al. Reactive glia not only associates with plaques but also parallels tangles in Alzheimer’s disease. Am J Pathol. 2011;179:1373–84.

9. Smit T, Deshayes NAC, Borchelt DR, Kamphuis W, Middeldorp J, Hol EM. Reactive astrocytes as treatment targets in Alzheimer’s disease-Systematic review of studies using the APPswePS1dE9 mouse model. Glia. 2021;69:1852–81.

10. Rodriguez-Vieitez E, Ni R, Gulyás B, Tóth M, Häggkvist J, Halldin C, et al. Astrocytosis precedes amyloid plaque deposition in Alzheimer APPswe transgenic mouse brain: a correlative positron emission tomography and in vitro imaging study. Eur J Nucl Med Mol Imaging. 2015;42:1119–32.

11. Ballweg A, Klaus C, Vogler L, Katzdobler S, Wind K, Zatcepin A, et al. [(18)F]F-DED PET imaging of reactive astrogliosis in neurodegenerative diseases: preclinical proof of concept and first-in-human data. J Neuroinflammation. 2023;20:68.

12. Olsen M, Aguilar X, Sehlin D, Fang XT, Antoni G, Erlandsson A, et al. Astroglial Responses to Amyloid-Beta Progression in a Mouse Model of Alzheimer’s Disease. Mol Imaging Biol. 2018;20:605–14.

13. Nam M-H, Ko HY, Kim D, Lee S, Park YM, Hyeon SJ, et al. Visualizing reactive astrocyte-neuron interaction in Alzheimer’s disease using 11C-acetate and 18F-FDG. Brain. 2023:awad037.

14. Kreimerman I, Reyes AL, Paolino A, Pardo T, Porcal W, Ibarra M, et al. Biological Assessment of a (18)F-Labeled Sulforhodamine 101 in a Mouse Model of Alzheimer’s Disease as a Potential Astrocytosis Marker. Front Neurosci. 2019;13:734.

15. Vilaplana E, Rodriguez-Vieitez E, Ferreira D, Montal V, Almkvist O, Wall A, et al. Cortical microstructural correlates of astrocytosis in autosomal-dominant Alzheimer disease. Neurology. 2020;94:e2026–e36.

16. Carter SF, Schöll M, Almkvist O, Wall A, Engler H, Långström B, et al. Evidence for astrocytosis in prodromal Alzheimer disease provided by 11C-deuterium-L-deprenyl: a multitracer PET paradigm combining 11C-Pittsburgh compound B and 18F-FDG. J Nucl Med. 2012;53:37–46.

17. Rodriguez-Vieitez E, Saint-Aubert L, Carter SF, Almkvist O, Farid K, Schöll M, et al. Diverging longitudinal changes in astrocytosis and amyloid PET in autosomal dominant Alzheimer’s disease. Brain. 2016;139:922–36.

18. Villemagne VL, Harada R, Doré V, Furumoto S, Mulligan R, Kudo Y, et al. First-in-Humans Evaluation of (18)F-SMBT-1, a Novel (18)F-Labeled Monoamine Oxidase-B PET Tracer for Imaging Reactive Astrogliosis. J Nucl Med. 2022;63:1551–9.

19. Villemagne VL, Harada R, Dore V, Furumoto S, Mulligan R, Kudo Y, et al. Assessing reactive astrogliosis with (18)F-SMBT-1 across the Alzheimer’s disease spectrum. J Nucl Med. 2022..

20. Chatterjee P, Pedrini S, Stoops E, Goozee K, Villemagne VL, Asih PR, et al. Plasma glial fibrillary acidic protein is elevated in cognitively normal older adults at risk of Alzheimer’s disease. Transl Psychiatry. 2021;11:27.

21. Villemagne VL, Harada R, Dore V, Furumoto S, Mulligan R, Kudo Y, et al. First-in-human evaluation of (18)F-SMBT-1, a novel (18)F-labeled MAO-B PET tracer for imaging reactive astrogliosis. J Nucl Med. 2022.

22. Harada R, Hayakawa Y, Ezura M, Lerdsirisuk P, Du Y, Ishikawa Y, et al. (18)F-SMBT-1: A Selective and Reversible PET Tracer for Monoamine Oxidase-B Imaging. J Nucl Med. 2021;62:253–8.

23. Oddo S, Caccamo A, Shepherd JD, Murphy MP, Golde TE, Kayed R, et al. Triple-transgenic model of Alzheimer’s disease with plaques and tangles: intracellular Abeta and synaptic dysfunction. Neuron. 2003;39:409–21.

24. Jankowsky JL, Fadale DJ, Anderson J, Xu GM, Gonzales V, Jenkins NA, et al. Mutant presenilins specifically elevate the levels of the 42 residue beta-amyloid peptide in vivo: evidence for augmentation of a 42- specific gamma secretase. Hum Mol Genet. 2004;13:159–70.

25. Hu W, Pan D, Wang Y, Bao W, Zuo C, Guan Y, et al. PET Imaging for Dynamically Monitoring Neuroinflammation in APP/PS1 Mouse Model Using [(18)F]DPA714. Front Neurosci. 2020;14:810.

26. Kawamura K, Hashimoto H, Furutsuka K, Ohkubo T, Fujishiro T, Togashi T, et al. Radiosynthesis and quality control testing of the tau imaging positron emission tomography tracer [(18) F]PM-PBB3 for clinical applications. J Labelled Comp Radiopharm. 2021;64:109–19.

27. Liu Y, Zhu L, Plössl K, Choi SR, Qiao H, Sun X, et al. Optimization of automated radiosynthesis of [18F]AV-45: a new PET imaging agent for Alzheimer’s disease. Nucl Med Biol. 2010;37:917–25.

28. Wilson A, Garcia A, Chestakova A, Kung H, Houle S. A rapid oneLstep radiosynthesis of the βLamyloid imaging radiotracer NLmethylL[11C]2L(4′Lmethylaminophenyl)L6Lhydroxybenzothiazole ([11C]L6LOHLBTAL1). J Labelled Comp Radiopharm. 2004;47:679–82.

29. Kong Y, Huang L, Li W, Liu X, Zhou Y, Liu C, et al. The Synaptic Vesicle Protein 2A Interacts With Key Pathogenic Factors in Alzheimer’s Disease: Implications for Treatment. Front cell dev biol. 2021;9:609908-.

30. Ni R, Villois A, Dean-Ben XL, Chen Z, Vaas M, Stavrakis S, et al. In-vitro and in-vivo characterization of CRANAD-2 for multi-spectral optoacoustic tomography and fluorescence imaging of amyloid-beta deposits in Alzheimer mice. Photoacoustics. 2021;23:100285..

31. Vagenknecht P, Luzgin A, Ono M, Ji B, Higuchi M, Noain D, et al. Non-invasive imaging of tau-targeted probe uptake by whole brain multi-spectral optoacoustic tomography. Eur J Nucl Med Mol Imaging. 2022.

32. Tagai K, Ono M, Kubota M, Kitamura S, Takahata K, Seki C, et al. High-Contrast In Vivo Imaging of Tau Pathologies in Alzheimer’s and Non-Alzheimer’s Disease Tauopathies. Neuron. 2021;109:42–58.e8..

33. López-Picón FR, Keller T, Bocancea D, Helin JS, Krzyczmonik A, Helin S, et al. Direct Comparison of [(18)F]F-DPA with [(18)F]DPA-714 and [(11)C]PBR28 for Neuroinflammation Imaging in the same Alzheimer’s Disease Model Mice and Healthy Controls. Mol Imaging Biol. 2022;24:157–66.

34. Galea E, Morrison W, Hudry E, Arbel-Ornath M, Bacskai BJ, Gómez-Isla T, et al. Topological analyses in APP/PS1 mice reveal that astrocytes do not migrate to amyloid-β plaques. Proc Natl Acad Sci U S A. 2015;112:15556–61.

35. Jo S, Yarishkin O, Hwang YJ, Chun YE, Park M, Woo DH, et al. GABA from reactive astrocytes impairs memory in mouse models of Alzheimer’s disease. Nat Med. 2014;20:886–96.

36. Park JH, Ju YH, Choi JW, Song HJ, Jang BK, Woo J, et al. Newly developed reversible MAO-B inhibitor circumvents the shortcomings of irreversible inhibitors in Alzheimer’s disease. Sci Adv. 2019;5:eaav0316.

37. Chun H, Im H, Kang YJ, Kim Y, Shin JH, Won W, et al. Severe reactive astrocytes precipitate pathological hallmarks of Alzheimer’s disease via H(2)O(2)(-) production. Nat Neurosci. 2020;23:1555–66.

38. Biechele G, Sebastian Monasor L, Wind K, Blume T, Parhizkar S, Arzberger T, et al. Glitter in the Darkness? Non-fibrillar β-amyloid Plaque Components Significantly Impact the β-amyloid PET Signal in Mouse Models of Alzheimer’s Disease. J Nucl Med. 2021.

39. Deleye S, Waldron AM, Verhaeghe J, Bottelbergs A, Wyffels L, Van Broeck B, et al. Evaluation of Small-Animal PET Outcome Measures to Detect Disease Modification Induced by BACE Inhibition in a Transgenic Mouse Model of Alzheimer Disease. J Nucl Med. 2017;58:1977–83.

40. Waldron AM, Wintmolders C, Bottelbergs A, Kelley JB, Schmidt ME, Stroobants S, et al. In vivo molecular neuroimaging of glucose utilization and its association with fibrillar amyloid-β load in aged APPPS1- 21 mice. Alzheimers Res Ther. 2015;7:76.

41. Poisnel G, Dhilly M, Moustié O, Delamare J, Abbas A, Guilloteau D, et al. PET imaging with [18F]AV-45 in an APP/PS1-21 murine model of amyloid plaque deposition. Neurobiol Aging. 2012;33:2561–71.

42. Ni R, Gillberg P-G, Bogdanovic N, Viitanen M, Myllykangas L, Nennesmo I, et al. Amyloid tracers binding sites in autosomal dominant and sporadic Alzheimer’s disease. Alzheimer’s & Dementia. 2017;13:419–30.

43. Ni R, Gillberg PG, Bergfors A, Marutle A, Nordberg A. Amyloid tracers detect multiple binding sites in Alzheimer’s disease brain tissue. Brain. 2013;136:2217–27.

44. Ni R, Chen Z, Deán-Ben XL, Voigt FF, Kirschenbaum D, Shi G, et al. Multiscale optical and optoacoustic imaging of amyloid-β deposits in mice. Nat Biomed Engineering. 2022.

45. Cao L, Kong Y, Ji B, Ren Y, Guan Y, Ni R. Positron Emission Tomography in Animal Models of Tauopathies. Front Aging Neurosci. 2021;13:761913. doi:10.3389/fnagi.2021.761913.

46. Barron AM, Ji B, Fujinaga M, Zhang MR, Suhara T, Sahara N, et al. In vivo positron emission tomography imaging of mitochondrial abnormalities in a mouse model of tauopathy. Neurobiol Aging. 2020;94:140–8.

47. Ishikawa A, Tokunaga M, Maeda J, Minamihisamatsu T, Shimojo M, Takuwa H, et al. In Vivo Visualization of Tau Accumulation, Microglial Activation, and Brain Atrophy in a Mouse Model of Tauopathy rTg4510. J Alzheimers Dis. 2018;61:1037–52.

48. Ni R, Ji B, Ono M, Sahara N, Zhang MR, Aoki I, et al. Comparative In Vitro and In Vivo Quantifications of Pathologic Tau Deposits and Their Association with Neurodegeneration in Tauopathy Mouse Models. J Nucl Med. 2018;59:960–6.

49. Ono M, Sahara N, Kumata K, Ji B, Ni R, Koga S, et al. Distinct binding of PET ligands PBB3 and AV- 1451 to tau fibril strains in neurodegenerative tauopathies. Brain. 2017;140:764–80.

50. Kimura T, Ono M, Seki C, Sampei K, Shimojo M, Kawamura K, et al. A quantitative in vivo imaging platform for tracking pathological tau depositions and resultant neuronal death in a mouse model. Eur J Nucl Med Mol Imaging. 2022.

51. Endepols H, Anglada-Huguet M, Mandelkow E, Schmidt Y, Krapf P, Zlatopolskiy BD, et al. Assessment of the In Vivo Relationship Between Cerebral Hypometabolism, Tau Deposition, TSPO Expression, and Synaptic Density in a Tauopathy Mouse Model: a Multi-tracer PET Study. Mol Neurobiol. 2022;59:3402–13.

52. Filip T, Mairinger S, Neddens J, Sauberer M, Flunkert S, Stanek J, et al. Characterization of an APP/tau rat model of Alzheimer’s disease by positron emission tomography and immunofluorescent labeling. Alzheimer’s Research & Therapy. 2021;13:175.

53. Chaney AM, Lopez-Picon FR, Serrière S, Wang R, Bochicchio D, Webb SD, et al. Prodromal neuroinflammatory, cholinergic and metabolite dysfunction detected by PET and MRS in the TgF344-AD transgenic rat model of AD: a collaborative multi-modal study. Theranostics. 2021;11:6644–67.

54. Metaxas A, Thygesen C, Kempf SJ, Anzalone M, Vaitheeswaran R, Petersen S, et al. Ageing and amyloidosis underlie the molecular and pathological alterations of tau in a mouse model of familial Alzheimer’s disease. Sci Rep. 2019;9:15758.

55. Ni R. Positron Emission Tomography in Animal Models of Alzheimer’s Disease Amyloidosis. Pharmaceuticals (Basel). 2021.

56. Kreisl WC, Kim MJ, Coughlin JM, Henter ID, Owen DR, Innis RB. PET imaging of neuroinflammation in neurological disorders. Lancet Neurol. 2020;19:940–50.

57. Zhou R, Ji B, Kong Y, Qin L, Ren W, Guan Y, et al. PET Imaging of Neuroinflammation in Alzheimer’s Disease. Front Immunol. 2021;12:739130.

58. Chauveau F, Van Camp N, Dollé F, Kuhnast B, Hinnen F, Damont A, et al. Comparative evaluation of the translocator protein radioligands 11C-DPA-713, 18F-DPA-714, and 11C-PK11195 in a rat model of acute neuroinflammation. J Nucl Med. 2009;50:468–76.

59. Zhou R, Ji B, Kong Y, Qin L, Ren W, Guan Y, et al. PET Imaging of Neuroinflammation in Alzheimer’s Disease. Front Immunol. 2021;12:3750.

60. Venneti S, Lopresti BJ, Wang G, Hamilton RL, Mathis CA, Klunk WE, et al. PK11195 labels activated microglia in Alzheimer’s disease and in vivo in a mouse model using PET. Neurobiol Aging. 2009;30:1217–26.

61. Liu B, Le KX, Park MA, Wang S, Belanger AP, Dubey S, et al. In Vivo Detection of Age- and Disease-Related Increases in Neuroinflammation by 18F-GE180 TSPO MicroPET Imaging in Wild-Type and Alzheimer’s Transgenic Mice. J Neurosci. 2015;35:15716–30.

62. Chaney A, Bauer M, Bochicchio D, Smigova A, Kassiou M, Davies KE, et al. Longitudinal investigation of neuroinflammation and metabolite profiles in the APP(swe) ×PS1(Δe9) transgenic mouse model of Alzheimer’s disease. J Neurochem. 2018;144:318–35.

63. Sérrière S, Tauber C, Vercouillie J, Mothes C, Pruckner C, Guilloteau D, et al. Amyloid load and translocator protein 18 kDa in APPswePS1-dE9 mice: a longitudinal study. Neurobiol Aging. 2015;36:1639–52.

64. Takkinen JS, López-Picón FR, Al Majidi R, Eskola O, Krzyczmonik A, Keller T, et al. Brain energy metabolism and neuroinflammation in ageing APP/PS1-21 mice using longitudinal (18)F-FDG and (18)F-DPA- 714 PET imaging. J Cereb Blood Flow Metab. 2017;37:2870–82.

65. Park BN, Kim JH, Lim TS, Park SH, Kim TG, Yoon BS, et al. Therapeutic effect of mesenchymal stem cells in an animal model of Alzheimer’s disease evaluated by β-amyloid positron emission tomography imaging. Aust N Z J Psychiatry. 2020;54:883–91.

66. Chen YA, Lu CH, Ke CC, Chiu SJ, Chang CW, Yang BH, et al. Evaluation of Class IIa Histone Deacetylases Expression and In Vivo Epigenetic Imaging in a Transgenic Mouse Model of Alzheimer’s Disease. Int J Mol Sci. 2021;22.

67. Chiquita S, Ribeiro M, Castelhano J, Oliveira F, Sereno J, Batista M, et al. A longitudinal multimodal in vivo molecular imaging study of the 3xTg-AD mouse model shows progressive early hippocampal and taurine loss. Hum Mol Genet. 2019;28:2174–88.

68. Tournier BB, Tsartsalis S, Rigaud D, Fossey C, Cailly T, Fabis F, et al. TSPO and amyloid deposits in sub-regions of the hippocampus in the 3xTgAD mouse model of Alzheimer’s disease. Neurobiol Dis. 2019;121:95–105.

69. Javonillo DI, Tran KM, Phan J, Hingco E, Kramár EA, da Cunha C, et al. Systematic Phenotyping and Characterization of the 3xTg-AD Mouse Model of Alzheimer’s Disease. Front Neurosci. 2021;15:785276.

70. Tang Z, Chen Z, Min G, Peng Y, Xiao Y, Ni R, et al. NRF2 deficiency promotes ferroptosis of astrocytes mediated by oxidative stress in Alzheimer’s disease. bioRxiv. 2023:2023.03.12.532248.

71. Jiwaji Z, Tiwari SS, Avilés-Reyes RX, Hooley M, Hampton D, Torvell M, et al. Reactive astrocytes acquire neuroprotective as well as deleterious signatures in response to Tau and Aß pathology. Nat Commun. 2022;13:135.

72. Bellaver B, Povala G, Ferreira PCL, Ferrari-Souza JP, Leffa DT, Lussier FZ, et al. Astrocyte reactivity influences amyloid-β effects on tau pathology in preclinical Alzheimer’s disease. Nat Med. 2023.

73. Liddelow SA, Guttenplan KA, Clarke LE, Bennett FC, Bohlen CJ, Schirmer L, et al. Neurotoxic reactive astrocytes are induced by activated microglia. Nature. 2017;541:481–7.

74. Xiang X, Wind K, Wiedemann T, Blume T, Shi Y, Briel N, et al. Microglial activation states drive glucose uptake and FDG-PET alterations in neurodegenerative diseases. Sci Transl Med. 2021;13:eabe5640.

75. Livingston NR, Calsolaro V, Hinz R, Nowell J, Raza S, Gentleman S, et al. Relationship between astrocyte reactivity, using novel 11C-BU99008 PET, and glucose metabolism, grey matter volume and amyloid load in cognitively impaired individuals. medRxiv. 2021:2021.08.10.21261690.

76. Hamelin L, Lagarde J, Dorothée G, Potier MC, Corlier F, Kuhnast B, et al. Distinct dynamic profiles of microglial activation are associated with progression of Alzheimer’s disease. Brain. 2018;141:1855–70.

77. De Bastiani MA, Bellaver B, Brum WS, Souza DG, Ferreira PCL, Rocha AS, et al. Hippocampal GFAP-positive astrocyte responses to amyloid and tau pathologies. Brain Behav Immun. 2023;110:175–84.

78. Furman JL, Sama DM, Gant JC, Beckett TL, Murphy MP, Bachstetter AD, et al. Targeting astrocytes ameliorates neurologic changes in a mouse model of Alzheimer’s disease. J Neurosci. 2012;32:16129–40.

